# Circulating Microparticles: Optimization and Standardization of Isolation Protocols and Reassessment of Their Characteristics and Functions

**DOI:** 10.1101/2023.05.04.539499

**Authors:** Chen Zhang, Jiajia Hu, Yifan Shi, Yang Feng, Zeyang Li, Tianyu Zhang, Liwen Hong, Zi Dong, Yiding Tang, Zhengting Wang, Guang Ning, Guorui Huang

## Abstract

Microparticles (MPs) are convenient for clinical diagnosis, and have functional roles in signal transduction. Although the importance of MPs is being increasingly recognized, the diversity of isolated protocols for MPs results in a heterogeneous population of their unknown origins, even expands to uncertain functions. Here we systematically studied the composition of MPs at different centrifugal speed intervals, and found that 3000g was a critical centrifugation speed in determining new MPs composition. The platelet-derived particles accounted for more than 80% under 3000g, while only about 20% in MPs obtained over 3000g. Furthermore, we found that the function of new MPs was significantly different from that of traditional ones, such as procoagulation activity, anti-inflammation and clinical diagnosis etc. Thus, our work optimized the method of MPs isolation, clarified some characteristics and physiological functions that should belong to platelets rather than MPs, which will derive new conceptual MPs for its composition and function.

## Introduction

Extracellular vesicles (EVs) can be divided into exosomes, microparticles (MPs) (or microvesicles) and apoptotic bodies according to size and synthesize ^1^. MPs are heterogeneous plasma membrane vesicles with diameters of 50 nm to 1000 nm generated by a variety of cells such as platelets, endothelial cells, leukocytes and erythrocytes etc., via the budding of the outer cell membrane during cell activation or apoptosis^2–4^. As an additional mechanism for intercellular communication, MPs allow cells to exchange proteins, lipids and genetic material^2^. Accordingly, there is a growing belief that MPs are involved in a variety of processes, including procoagulant and fibrinolytic activities^5^, vascular remodeling or neo angiogenesis^6^, promoting cellular interactions and signal transmission^7^, and providing malignant cell microenvironments ^8^. However, the methods that are currently applied to isolate MPs are far from standardization, few of which can be directly compared to the others, and of which results are sometimes inconsistent or conflicting^9–13^. The method of MPs isolation generally consists of several steps, including a low-speed centrifugation to remove cells, platelets and apoptotic debris, and a high-speed centrifugation to precipitate MPs^14^. The common centrifugation parameters used for MPs isolation vary from 1000 to 5,000 g for 5–20 min in the initial centrifugation step, followed by 13,000–100,000g for 30–60 min to pellet MPs^9^. Therefore, the characteristics of MPs depend on the detection protocols because there is no uniform consensus regarding the definition of MPs ^15^. To avoid falsely high or low quantification of the composition and understand the real function of MPs, optimization and standardization of detection methods are of great importance.

Platelets are active participants in the immune response to microbial organisms and foreign substances by triggering platelet-neutrophil interactions^16, 17^. As revealed by recent studies, platelet activation also leads to circulating MP formation in addition to the well-known hemostatic and inflammatory responses described, which is becoming an increasing focus in medical research^18–20^. MPs were first described as subcellular material originating from platelets in normal plasma and serum and called ‘platelet dust’ formerly ^20^. It has long been thought that 70-90% of MPs in plasma are derived from platelets^14, 21, 22^. And it also has been studied mainly for their role in blood coagulation^23^, vascular remodeling or angiogenesis^6^, all of which are similar to the function of platelets. Therefore, according to the current mainstream MPs detection protocols, the vast majority of MPs would be derived from platelets, and thus their functions also depend on platelets. This obscures the characteristics and functions of many other cell-derived MPs, and also interferes with the results and conclusion about MPs studies.

In addition, high MP levels have been reported in diseases association and showed their diagnostic and prognostic value ^24–27^. Despite several decades of extensive studies, they have not shown clinical benefits from these MPs alterations in patients, because many other cell-derived MPs were blocked by the vast majority of “Platelet-derived MPs”. Here we found that the proportion of MPs derived from different cells varies greatly with different centrifugal speeds, especially at lower speeds. Then we systematically studied the composition of MPs in different centrifugal speed intervals, and found that after 3000g, whether 5000g, 8000g, 12,000g or 20,000g all were similar to each other, which means that the proportion of MPs from different cell sources is relatively stable. It indicated that the MPs obtained after 3000g may be special and even true MPs. After that, we further confirmed that the proportion of MPs derived from each cell was very stable after 3000g by electron microscopy, dynamic light scattering (DLS), immunofluorescence combined with flow cytometry et al. Finally, we reevaluated and found that some functions belong to platelets rather than MPs like the procoagulation activity, some belong to both of platelets and MPs like anti-inflammatory effect (but MPs better), some profit from other compositions of MPs like the promotion of cell migration and tube formation, and some used for clinical diagnosis such as cardiovascular disease (endothelial cell-derived MPs). Thus, our work established MPs isolation method and obtained true MPs, which will contribute to the future studies of its characteristics and physiological function.

## Results

### Diverse and confusing microparticle isolation methods

Recent works demonstrate MPs contain mitochondrial components as well as intact mitochondrial, which was described as mediators of inflammation ^28, 29^. Thus, the mitochondrial components from different cells were detected, and we found the proportion of mitochondrial characteristic MPs from different cells was almost similar. Actually, the MPs derived from platelets (CD41^+^) were reported with dozens of times higher than those from other cell sources (**Figure S1**). This made us have to suspect that the currently MPs isolation methods might have potential problems, and many of MPs may be the fragments of platelets rather than true MPs. Although MPs have been extensively studied and found to be diverse in their functions, there is a lack of accepted uniform standard for their isolation methods (**Table S1**). In particular, the initial centrifugation speed for removing cell fragment is rarely the same, which results in a large difference in its composition, but most of them are platelet-derived and account for 70-90%. The diversity of components naturally leads to the diversity of their functions, and some of them are even contradictive (**Table S1)**. For example, it has been reported to promote tumor metastasis^30^, but it has also been reported to inhibit cell migration and proliferation on the contrary ^31^. Of course, the most studies of its functions are that it promotes coagulation and angiogenesis, and inhibits inflammatory responses. Because most published studies of MPs have focused on their potential functions rather than on their origins, it is still unclear which cell-derived particles is responsible for any given function^2^. To clarify these confusions and uncertainties about MPs studies, it is necessary to systematically study the composition of MPs at different centrifugation speeds and redefine their physiological functions.

**Table SI.**
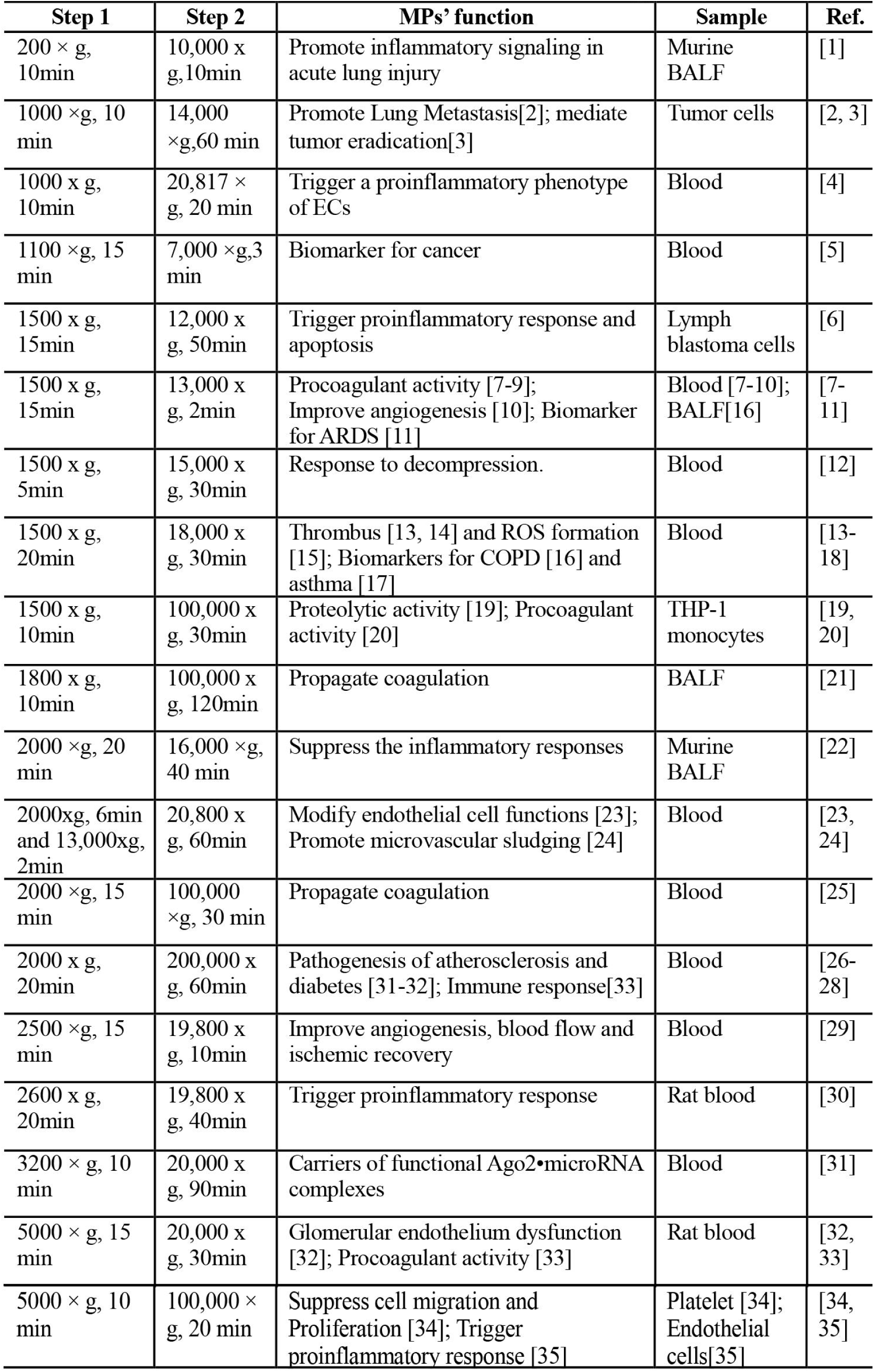
Characteristics of MPs with different published centrifugation protocols.

**Figure S1.**
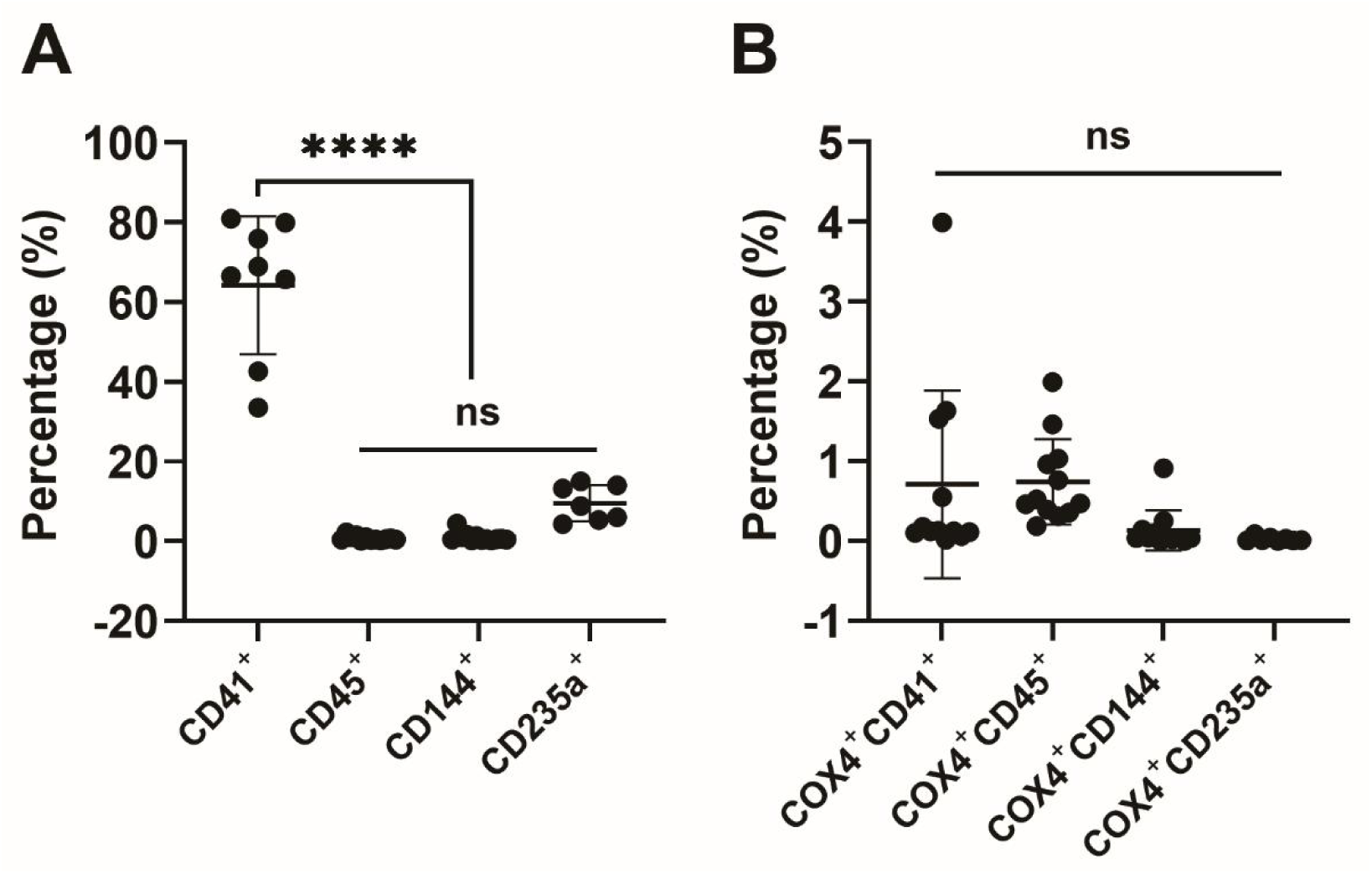
Detection of circulating cell-derived MPs (A) or with mitochondrial positive (Cox4^+^) (B).

### The composition of MPs obtained at different centrifugal speed intervals

Although the initial centrifugal speed diversified from different literatures, most of them centrifuged at around 1500g for 20 min to remove the cells and cell debris, and then harvested the microparticle pellets at around 20,000g for 30 min. To investigate the accurate centrifugal speed for detecting the stable MPs, particles were isolated at 20,000g after different initial concentrations like: 1500g, 2000g, 2500g, 3000g, 3500g and 4000g-post, respectively. Data showed that the proportion of CD41^+^ decreased according to the higher centrifugal speeds, and it became smoothy stable after 3000g-post (**Figure 1A, B**). It indicated that the 3000g may be the critical centrifugal speed for MPs isolation. Furthermore, we isolated particle samples at different centrifugal speeds intervals after initial 1500g as: 3000g, 5000g, 8000g, 12,000g and 20,000g. Consistently, over 80% particles are CD41^+^ under 3000g centrifugation. However, only ∼20% CD41^+^ MPs were found at all of other centrifugal speed intervals, which expressed pretty stable proportion (**Figure 1C**). It indicated that centrifugation at 3000 g for 20 min have eliminated small platelet and platelet fragmentation, and the rest of CD41^+^ MPs seemed to be the real platelet-derived-MPs. In contrast, compared with only 0.26% of the CD45^+^ (leukocyte-derived) MPs obtained at 3000g, the percent of CD45^+^ MPs isolated from other centrifugal speed intervals after 3000g were ∼2.2%, which was ∼10 times higher, and remained relatively stable (**Figure 1D**). Likewise, endothelial-derived MPs (CD144^+^) obtained at 5000g and other higher centrifugation speeds (∼3%) were approximately 6 times higher than 3000g (∼0.5%), and were also very stable (**Figure 1E**). More importantly, because the presence of a large number of particles were not true MPs, the proportion of mitochondria-positive MPs harvested at 3000g was very low, while it was around 3-7% when centrifuged at higher than 3000g centrifugation speed, which was much higher than that obtained at 3000g (**Figure 1F**). These data all indicated that 3000g was a critical centrifugation speed in determining MPs composition. Interestingly, the erythrocyte-derived MPs (CD235a^+^) obtained at 5000g were only slightly higher than those obtained at 3000g, but not significant different (**Figure 1G**). However, obtaining CD235a^+^ MPs was much higher when centrifugal over 5000g. Considering that the proportion of other cell-derived MPs was relatively consistent when centrifuged at over 3000g, and was significantly different from that obtained at 3000g. Meanwhile, particle samples under 3000g were detected a 10-fold higher amount of protein concentration compared to the other groups, which means that the most of the microparticle proteins obtained after 1500g was that of 3000g (**Figure 1H**). These findings suggested that when intact cells and platelets were removed at 1500g for 20 min, the directly collected MPs contained mainly platelet fragments or small platelets, as well as some erythrocyte debris, instead of particles that were not produced by cellular physiological processes. The debris can be largely removed after another 20 minutes of centrifugation at 3000g, and the plasma particles obtained are the stable and real cells-derived MPs.

**Figure 1.**
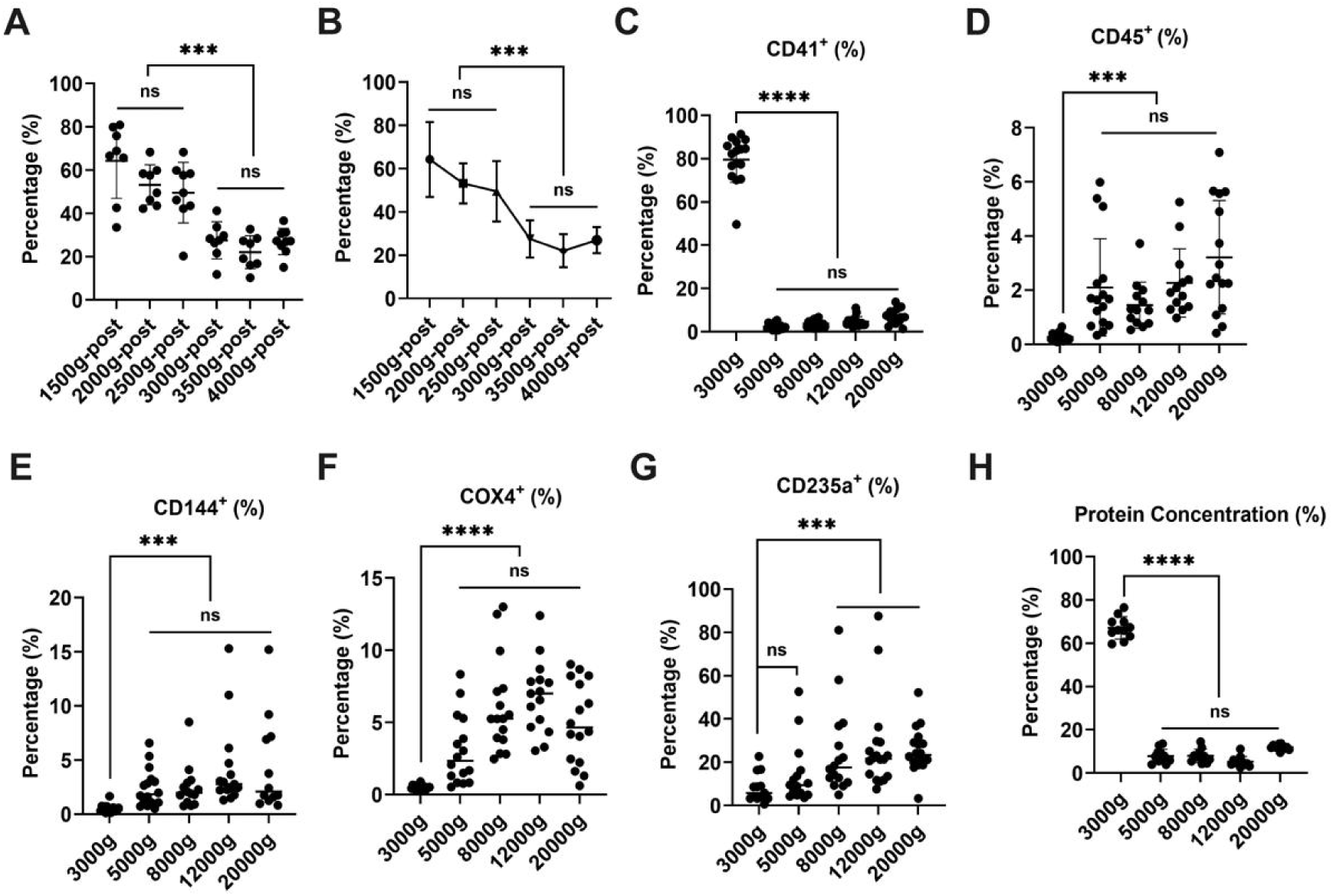
Detection of circulating cell-derived MPs at centrifugal speed intervals. (**A-B**) The percentage of CD41^+^ were detected when the particles were harvested at 20,000g after initial centrifugation of 1500g, 2000g, 2500g, 3000g, 3500g and 4000g, respectively. **(C)** After being centrifugated under different speeds, the percentage of platelet-derived (CD41^+^), leukocyte-derived (CD45^+^) (**D**), vascular endothelial cells-derived (CD144^+^) (**E**), mitochondrial-derived (COX4^+^) (**F**) and erythrocyte-derived (CD235a^+^) MPs (**G**) were shown, respectively. (**H**) The percentage of proteins obtained at five different centrifugation speeds relative to total particles proteins. n = 8-10 per group, data are presented as mean ± SD. *** P < 0.001, **** P < 0.0001.

### Analyze MPs by combining with immunofluorescence and flow cytometry

To further validate above results of flow cytometry, we combined immunofluorescence and flow cytometry to investigate the MP fractions in plasma obtained at different centrifugation speeds. The MPs were labeled with the detection antibodies for the cellular markers, and the secondary antibodies used red fluorescence. Consistent with the above results, direct observation by confocal microscopy showed that platelet-derived MPs obtained at 3000 g after 1500 g accounted for the majority of the MPs (**Figure S2A**), significantly higher than those from other cell sources such as leukocytes and erythrocytes (**Figure S2A-C**). The same samples were also quantitatively analyzed by flow cytometry again, and results were similar to those in above **Figure 1** (**Figure S2D**). In addition, western blotting was used to further confirm the relevant results, and found that the bands of platelet marker proteins CD41 and CD36 in 3000g-sample were much stronger than those obtained at other centrifugation speeds (**Figure S2E**). Importantly, the markers of other cells such as erythrocytes (glycophorin), leukocytes (CD45) and membrane lipoprotein (Perilipin1) obtained at different speeds did not differ significantly. However, as a cytoskeletal protein marker in cytosol, the band of Actin in 3000g-sample was much stronger than that of other centrifugation speeds, which was similar to platelet marker CD41 and CD36 (**Figure S2E**). It further suggested that most of the 3000g-particles were big cellular fragments or debris, and a small number of erythrocyte fragments, rather than the MPs produced by physiological processes.

**Figure S2.**
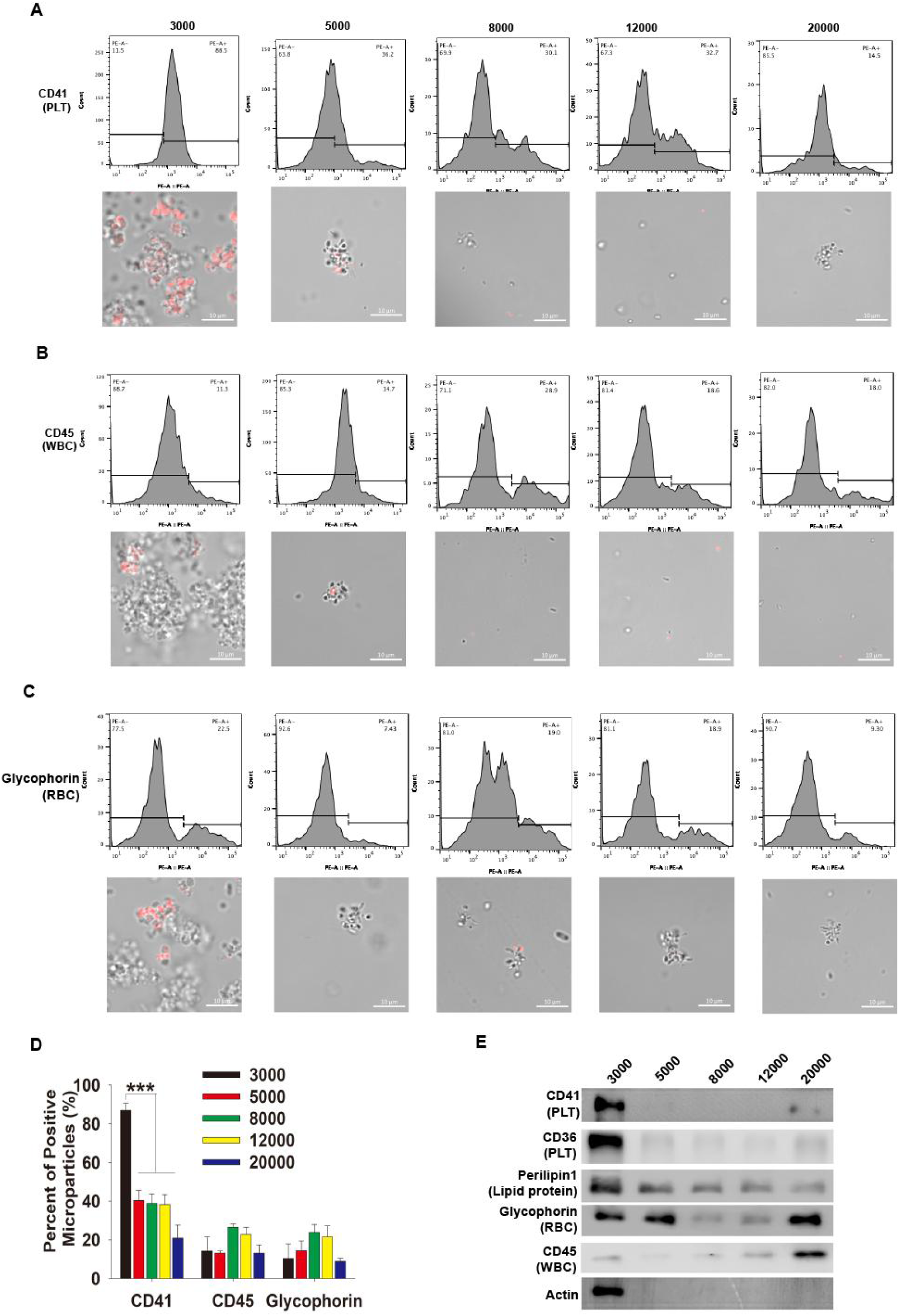
Validation of circulating MPs origins at different centrifugal intervals. (**A**) Representative images of immunofluorescent staining and flow cytometry analysis for platelet-derived (CD41^+^), leukocyte-derived (CD45^+^) (**B**) and erythrocyte-derived particles (Glycophorin) (**C**) at indicated five different centrifugation speeds. (**D**) Statistical analysis of CD41, CD45 and glycophorin positive MPs at indicated centrifugal speeds. (**E**) Western blotting analysis of particles were isolated under different centrifugation speeds with antibodies against CD41, CD36, Perilipin1, Glycophorin, CD45 and Actin. Data are mean ± SD; n = 4 per group. *** P < 0.001.

### The proportional MPs with centrifugal speeds

Although the initial centrifugal speed diversified from different literatures, most of them centrifuged at around 1500g for 20 min to remove the cells and cell debris, and then harvested the MPs pellet at around 20,000g for 30 min. To detect how the initial centrifugal speed determine the proportions of MPs, we compared the 1500g (1500g-post) and 3000g (3000g-post) as the initial speed to remove cell debris, respectively. As shown in **Figure 2A**, the protocols for the isolation of circulating particles were summarized. As expected, over 70% pellets were platelet-derived (CD41^+^) for 1500g-post as previous reports (**Figure 2B**). However, only ∼20% platelet-derived MPs were found when they were harvested at 3000-post, which is 2-3 times less than 1500g-post. Conversely, all of other cell-derived MPs like leukocyte (CD45^+^) (**Figure 2C**), vascular endothelial cells (CD144^+^) (**Figure 2D**) and mitochondrial-derived MPs (Cox4^+^) (**Figure 2E**) increased over 5 folds in 3000g-post compared with 1500g-post group. Interestingly, no significant increase was found in 3000g-post erythrocyte-derived MPs (CD235a^+^) (**Figure 2F**), most likely because there was still some of erythrocyte-derived debris present in the 1500g-post particles. Besides, the total proteins of MPs obtained at 3000g were detected around 4-fold amount of 3000g-post, it also means that the amount of microparticle proteins obtained from 3000g-post was only ∼20% of that of 1500g-post, and∼80% of the proteins in 1500g-post could be removed at 3000g (**Figure 2G**). It indicated that most of the so-called “MPs” obtained by1500g-post are mainly derived from small platelets and big platelet debris, and some of them also derive from erythrocyte fragments. These results further demonstrated that the centrifugal speed does play a critical role in MPs’ compositions, and 3000g is the boundary centrifuged speed for MPs isolation.

**Figure 2.** The percent of the MPs compositions and the amount of protein according to 1500g-post and 3000g-post centrifugation. (**A**) The summarized protocols for the isolation of circulating particles. (**B**) The percentage of platelet-derived (CD41^+^), leukocyte-derived (CD45^+^) (**C**), vascular endothelial cells-derived (CD144^+^) (**D**), mitochondrial-derived (COX4^+^) (**E**), and erythrocyte-derived MPs (CD235a^+^) (**F**) were shown, which obtained at 1500g-post and 3000g-post. (**G**) The percentage of protein after being centrifuged at 3000g and then 20000g (3000g-post) one after another, the total protein is equal to the protein of 1500g-post. Data are mean ±SD; n > 10 per group. *** P < 0.001, **** P < 0.0001.

### Morphological characterization and size of MPs

Size is a basic characteristic of MPs, and the size of MPs is recognized about 50-1000 nm, whose range is very wide^2, 3^. In order to further understand the size of MPs obtained by 3000g-post, we firstly used DLS to detect the size distribution of MPs obtained by 1500g-post (traditional), 3000g and 3000g-post (modified), respectively (**Figure 2A**). The data showed that the MPs obtained by the traditional method obviously had two peaks (**Figure 3A**). In terms of the proportion of number, the main peak is mainly at 150-500 nm (∼85%), another weaker main peak was around 600-1600 nm (∼15%) (**Figure 3A**), but it took up ∼80% amount of the total protein (**Figure 2G**). Interestingly, the “MPs” obtained at 1500g and then at a relative low speed of 3000g showed multiple peaks, especially the “MPs” over 1000 nm showed a plateau peak of ∼5% until over 2000 nm particles (total ∼48.9%) (**Figure 3B**). Although there was also a peak in the 100-500 nm region of 3000g particles, the total number proportion was only ∼40%. There was also a peak at 500-800nm, which accounted for about ∼11% (**Figure 3B**). What was more interesting was that the MPs obtained after our improved separation method had only one main peak at 100-500 nm, and its number accounted for 99% (**Figure 3C**). Additionally, the superposition of the size distribution of particles by three different methods also showed obvious differences (**Figure 3D**). All of these showed that the size of MPs obtained by the modified protocol has good uniformity and unique distribution, suggesting that our improved method is excellent for the definition of MPs without causing confusion, generality and uncertainty.

**Figure 3.**
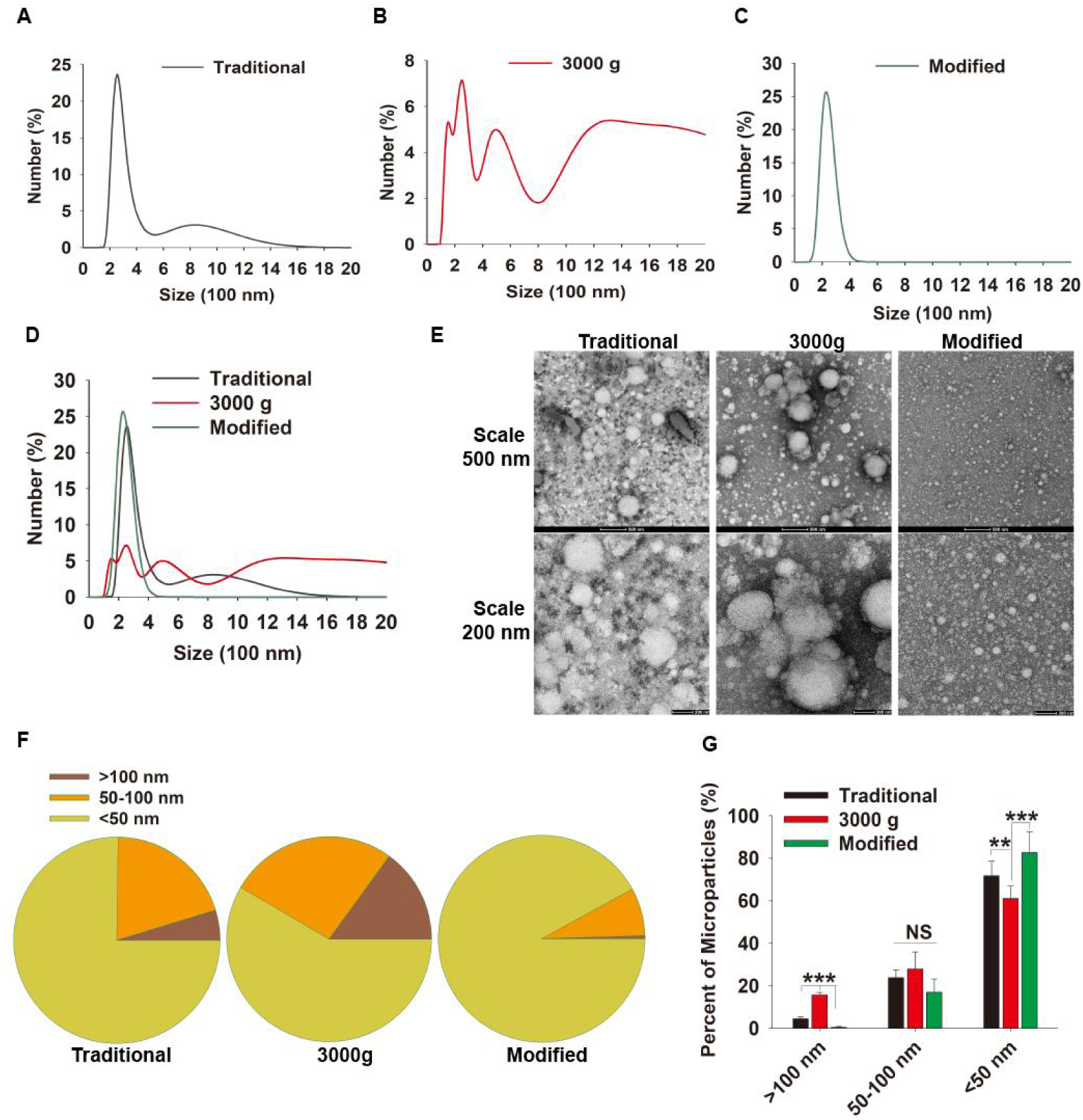
Morphological characteristics and size of MPs. (**A**) The size distribution of MPs was analyzed by the dynamic light scattering (DLS) for samples which obtained with traditional (1500g-post), 3000g (**B**) and modified (3000g-post) (**C**) protocols. (**D**) The three sizes distribution of MPs were merged. (**E**) Representative images were shown for particles morphological analysis by negative stain transmission electron microscopy (TEM). (**F**) Statistic analyses for the proportion of different sizes of particles. (**G**) Percentages of MPs were quantified for different size distributions from three different methods. Data are mean ±SD; over 10 images per group were analyzed. ** P < 0.01, *** P < 0.001.

To observe the particles more intuitively, we used transmission electron microscopy (TEM) with negative staining to analyze their morphology and size. Consistent with the DLS, the MPs obtained by the traditional method had both large and small particles, while the “microparticle” obtained by 3000g had greatly reduced small particles (**Figure 3E**). However, the MPs obtained by modified method showed pretty uniformity and unique (**Figure 3E**). Various statistical analyses had shown significant differences from traditional methods (**Figure 3F, G**). It should be pointed out that the size of the MPs observed by TEM was smaller than that detected by the DLS method. We speculated that this was mainly because the MPs needed to be fixed during the preparation of the electron microscope samples, then the MPs were negatively stained, and dried with heat lamp. All of them resulted in the shrinkage of plasma particles and a significant reduction in the observed sample size due to the membrane structure of particles. And this speculation confirmed by DLS data, which showed the size of samples much smaller after being prepared as above TEM methods (**Figure S3**). Thus, the actual size of which should be determined by the DLS method without fixation, and the particles were resuspended in solution. In view of this, the size of new MPs should be within the 100-500 nm range, and the vast majority are distributed in a narrower range between 200-350 nm (∼85%).

**Figure S3.**
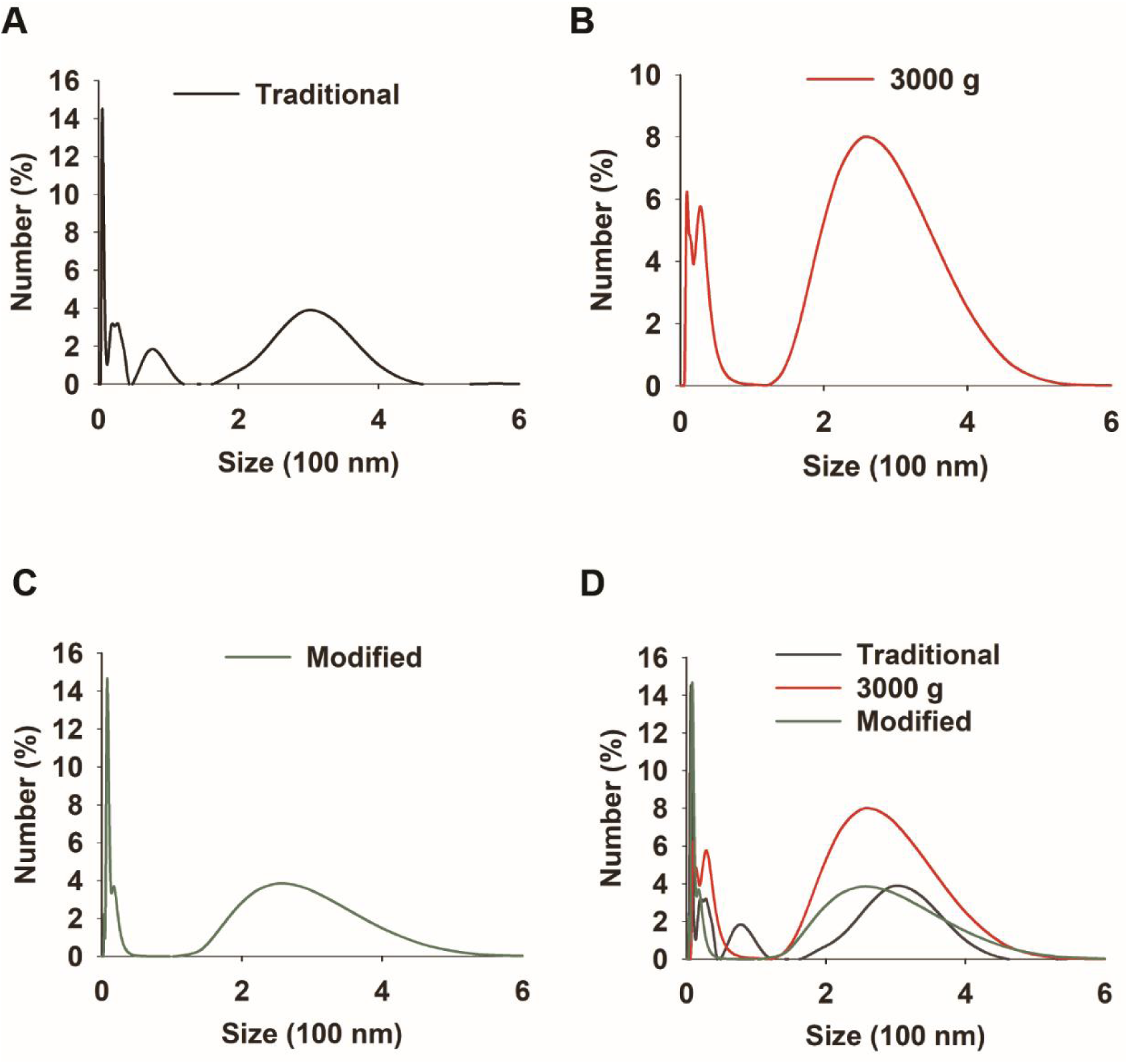
The size distribution of MPs was analyzed by the dynamic light scattering (DLS) for indicated samples after being prepared with TEM methods.

### Biochemical detection of cells-derived MPs

To further detect the characteristics of MPs with different methods, we used biochemical methods like immunofluorescence combined with flow cytometry and western blotting. As mentioned above, the particles isolated by traditional method (1500g-post) were similar to that obtained by centrifugation at 3000g, and the proportion of CD41-positive particles reached more than 80%, which was much higher than that of our improved method (∼30%) (**Figure 4A**). Besides, the particles obtained at 1500g-post and 3000g were more numerous and existed in agglomerate, rather than smaller and dispersed as in 3000g-post MPs (**Figure 4A-C**). Similarly, for leukocyte- and erythrocyte-derived MPs, both the traditional and modified methods did not differ significantly, but they all had significantly higher proportion of positive particles obtained at 3000g (**Figure 4D**). In addition, we detected various cell-derived proteins (derived from an equal volume of plasma, of which modified and 3000g groups were derived from the same sample, and the samples preparation below is the same) by Western blotting. It was found that the platelet-derived protein (CD31) in 3000g-post samples was significantly lower than that in 1500g-post and 3000g. However, the other several marker proteins like Perllipin1 (lipid protein), Glycophorin (RBC) and CD45 (WBC) were not much different, they were all significantly more than what the 3000g sample obtained (**Figure 4E**). More importantly, as the cytoskeletal protein marker existing in the cytoplasm, whether in the 1500g-post or 3000g group, the expression of Actin was much higher than that obtained by our improved method, which further indicated that the particles in 1500g-post and 3000g mainly came from cell fragmental particles rather than plasma membrane-dominant vesicles (**Figure 4E**). Consistent with the previous results, these data demonstrated that the majority of proteins isolated by traditional methods were platelet-derived particles and platelet fragments rather than plasma membrane vesicle MPs.

**Figure 4.**
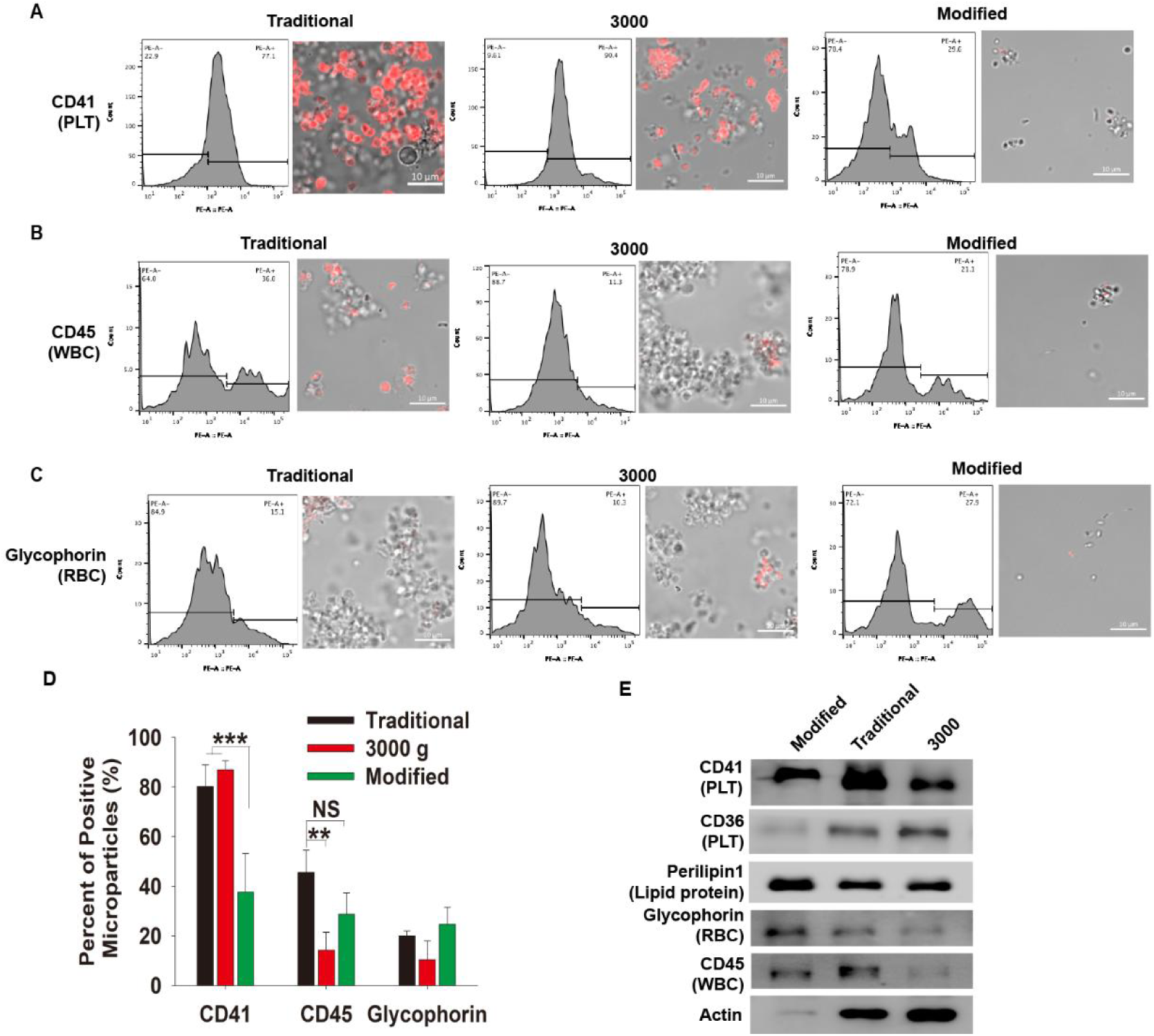
Composition of isolated MPs with different centrifugations. (**A**) Immunofluorescent staining and flow cytometer analyze for the MPs from different cell-derived markers like CD41, CD45 (**B**) and Glycophorin (**C**) under traditional, 3000g and modified centrifugation protocols. (**D**) Statistical analysis of the percentage of CD41, CD45 and Glycophorin positive MPs at different speeds. (**E**) Western blotting analysis for particles isolated under indicated speeds with antibodies against CD36, CD45, Glycophorin, Perillipin1, and cytoskeleton protein Actin. Data are mean ± SD; n = 4 per group. ** P < 0.01, *** P < 0.001.

### Procoagulant function belongs to platelet granules rather than MPs

Procoagulant activity and thrombus formation are well-known important function of MPs^5, 22^. To validate the procoagulant activities and thrombus formation, MPs isolated by different centrifugation protocols were added into the general cell culture medium DMEM with 10% FBS. A clot was quickly formed in the medium that contains particles obtained at traditional centrifugation methods (1500g-post) or 3000g. However, the medium had been kept clarifying when it was added the MPs harvested with modified protocol (3000g-post) (**Figure 5A**). Furthermore, the impact of MPs on procoagulant function and microthrombosis was evaluated via *in vitro* coagulation tests. As shown in **Figure 5B** and **5C**, obvious coagulation was observed at ∼60-90s in 3000g and 1500g-Post treatments, but no significant differences were found between control and 3000g-Post. This data suggested that the procoagulant function of 1500g-post “MPs” in many literatures is largely dependent on the presence of the large number of platelets and platelet debris, which can be isolated at 3000g, rather than the true function of plasma membrane vesicle MPs that require higher speed centrifugation. Thus, the procoagulant activities of real MPs need to be reevaluated.

**Figure 5.** The Roles of MPs in Procoagulant activity and as Biomarker of Endothelial Damage in CAD. (**A**) Normal cell cultured DMEM medium supplement with 10% FBS were treated with the indicated particles for 2 min, and then the clots were pick up with forceps. (**B**) *In vitro* coagulation test and clotting time **(C)** in different MPs groups. **(D)** CD144^+^ MPs were analyzed by flow cytometer for samples which obtained with 1500g-post, 3000g and 3000g-post protocols. The difference of CD144^+^ MPs percentage between CAD and healthy individuals was shown for samples obtained with 1500g-post protocols **(E)** and 3000g-post **(F)** protocols. Data are mean ± SD. ** P < 0.01, **** P < 0.0001.

### Endothelial derived MPs (EMPs) reflect coronary artery disease (CAD)

CAD is a major cause of mortality and morbidity across world and showed pro-inflammatory and pro-coagulant condition of body. The present findings support the view that the CD31^+^ EMP is a sensitive marker for CAD based on data from some groups ^32, 33^. Although the levels of CD144^+^ EMPs increased in patients with CAD and predict CAD more strongly than traditional risk factors ^34, 35^. The correlation between CD144^+^ and CAD was very poor and no significant difference was found ^36^. In order to detect the difference between traditional and our modified MPs isolation method in diagnosis of diseases, the CD144^+^ EMPs in CAD were studied firstly. 93 patients with CAD scheduled for invasive angiography and percutaneous coronary intervention (PCI) were included in the study, patients at a physical examination center were recruited as a comparator group. As expected, the level of CD144^+^ EMPs harvested with modified protocol was ∼5 folds higher than with the traditional protocol and 3000g (**Figure 5D**). Interestingly, the CAD patients had 30-40% higher levels of CD144^+^ EMPs, compared to the healthy control in 3000g-post group (**Figure 5E**), while no significant difference was found in 1500g-post EMP (CD144^+^) (**Figure 5F**). These results suggested that the new microparticles might predict enhanced vascular injury in individuals with CAD as biomarker and provide a novel therapeutic target in CAD.

### Promote angiogenesis and cell migration

MPs are reported to exhibit proangiogenic activity by promoting formation of capillary-like structures^6, 37^. To investigate the angiogenic effect of MPs, we performed a tube formation assay with HUVEC (Human Umbilical Vein Endothelial Cells) cells cultured in Matrigel in the presence of MPs isolated by different protocols. The formation of tubular structures was imaged after being cultured on Matrigel for 2 hours (**Figure 6A**). Interestingly, compared to that of in control and 3000g groups, the tubules were denser, and some tubules were darker in both traditional and modified protocol groups, and 3000g-post seemed more evident (**Figure 6A**). Following imaging of the complexes, we analyzed the characteristic information of the networks to quantify and compare the angiogenesis potentiality among the groups. Consistent with previous reports, groups treated with MPs isolated with modified and traditional methods, had more evolved networks in terms of number of nodes and total length (**Figure 6B, C**). And the formation of tube treated with the MPs from modified method was the best of all, but the characteristics of tubes treated with 3000g particles is similar to that of non-treated control (**Figure 6B, C**). These results suggested that the new MPs promote angiogenesis indeed, but platelet-derived fragments are not or much weaker. Thus, it indicated that previously reported the pro-angiogenic function is the function of modified MPs itself.

**Figure 6.**
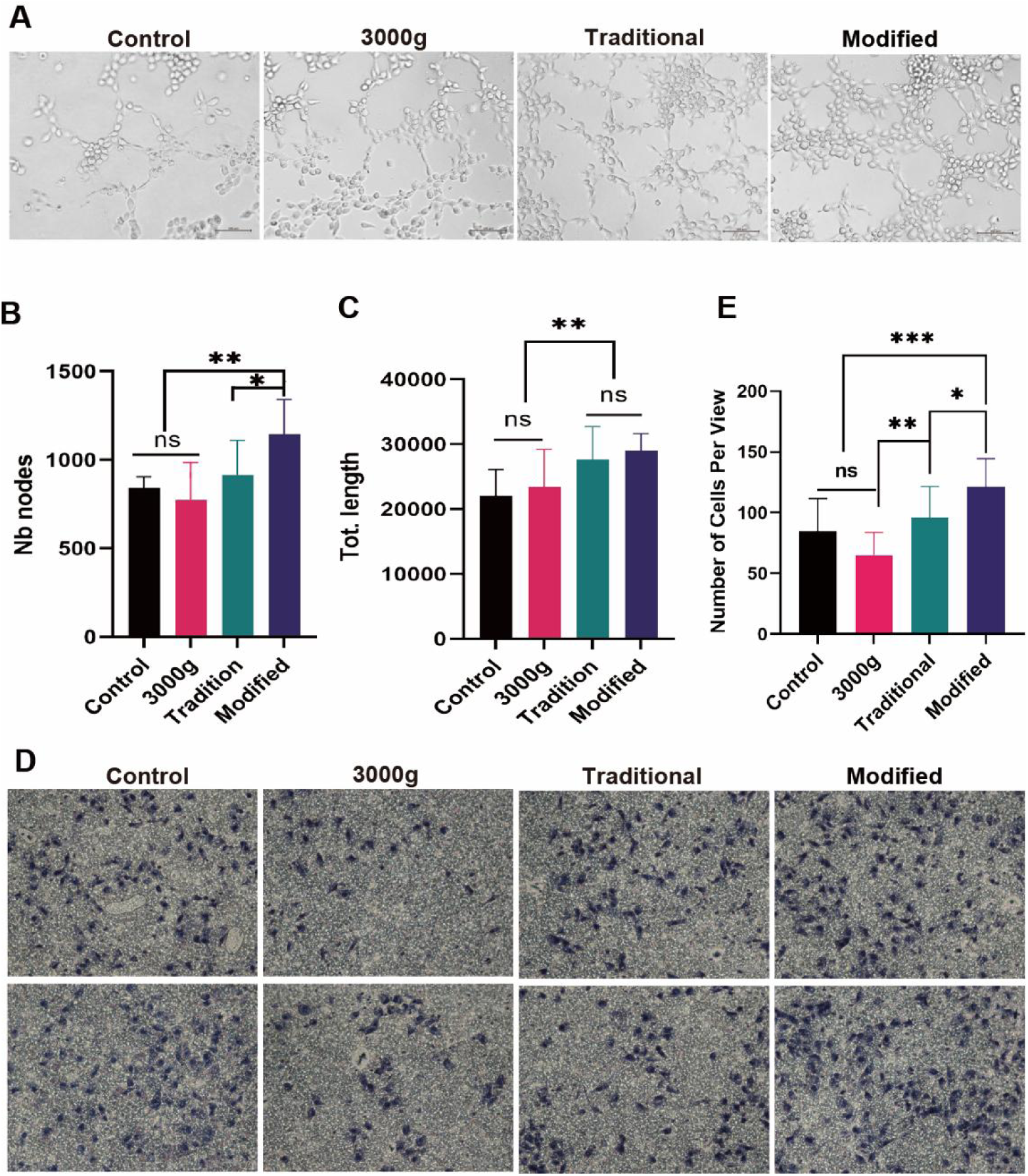
The effects of MPs on angiogenesis and cell migration. (**A**) Tube formation assay was performed in Matrigel and treated with indicated particles for 2 hours, and representative images of the endothelial tubular networks were shown. (**B**) Quantification of the network characteristics like the number of tube nodes and total length of tubes (**C**) with Image J. (**D**) For transwell migration assay HUVEC cells were treated with traditional, 3000g and modified MPs for 24 h, which imaged with 5x magnification and quantified with Image J (**E**). Data are mean ± SD; n = 5 per group. * P < 0.1, ** P < 0.01, *** P < 0.001.

Some of studies have reported that MPs could suppress cell migration^31, 38^. Thus, the *in vitro* transmigration assay was also performed to investigate the effects of MPs on the cell migration. Our results demonstrated that MPs obtained from both modified and traditional methods increased the HUVEC cells migration, but not for 3000g sample (**Figure 6D, E**). Thus, similar to promoting angiogenesis, the function of promoting cell migration may more likely belong to other cell-derived MPs, rather than platelet-derived particles.

### Suppress the inflammatory responses

Although Garnier, Y., et al., reported that MPs isolated from sickle patient plasma could trigger a proinflammatory phenotype of ECs^39^. Most of other studies found that the MPs suppress the inflammatory responses ^19–21, 40^. In order to determine the effect of the particles obtained by different methods on the inflammatory response, we first stimulated Mice Immortalized Bone Marrow-Derived Macrophage (iBMDM) with lipopolysaccharide (LPS) to develop an inflammatory model *in vitro* and treated with MPs. Interestedly, we found that after MPs treatment, the RNA levels of inflammatory factors IL-1β and its regulated protein NLRP3 were significantly decreased, much less than the positive control LPS stimulation (**Figure 7A, B**). The results showed that the protein level of IL-1β increased after MPs treatment, but only its precursor form increased, and its functional form of mature IL-1β was significantly lower than the control group and LPS treatment group, which may due to the decrease of NLRP3 (**Figure 7C**).

**Figure 7.** The effect of MPs on inflammatory response. (**A-C**) iBMDM cells were treated at indicated treatment for 6 h, and then harvested the samples and the mRNA level of NLRP3 **(A)** and IL-1β **(Β)** were detected, and Western blotting to detect indicated proteins **(C)**. (**D**) ALI mouse model obtained by being instilled i.t. LPS, isolated the MPs from the plasma of these mice 1 day later, and then instilled i.t. other mice together with LPS. 1 day later, BALF was harvested with 700 ul PBS. (**E**) Quantified the proteins in BALF from different treatments. (**F**) The cells from BALF were harvested and the levels of IL-1β and NLRP3 were detected with Western blotting. (**G**) The acute inflammatory regions in lung stained with H&E after being instilled i.t. LPS and treated with indicated MPs, and their quantification (**H**).

In addition, in order to further detect how MPs affected the inflammatory response, we used LPS to induce acute lung injury (ALI) mouse model for inflammation, and 1 day later, instilled i.t. with particles obtained at 3000g, 1500g-post and 3000g-post to observe the effect of MPs on the inflammatory response. Compared with control mice, the color of bronchoalveolar lavage fluid (BALF) of the mice injected with MPs became significantly lighter, indicating that the hemoglobin content of them was greatly reduced, and the traditional and modified methods of higher speed centrifugation were especially more effective (**Figure 7D**). The protein concentration in the BALF was detected, and it was also found that the protein content of the groups injected with MPs or cell fragment was significantly lower than that of the control group (**Figure 7E**). Furthermore, data showed that the protein level of IL-1β increased after MPs treatment, but only its precursor form increased, and its functional form of mature IL-1β was significantly lower than the control group, which may due to the decrease of NLRP3 (**Figure 7F**). Importantly, the inflammatory response in injury lung in MPs treatment have been improved than that in control group, it showed that all mice instilled i.t. with MPs had less infiltrations of immune cells and improved endothelial permeability, especially the 3000g-post obtained MPs treatment had a very significant anti-inflammation effect. (**Figure 7G, H**). Therefore, the results demonstrated that both MPs and platelet-derived particles can significantly suppress the inflammatory response in cells and mice. And the modified MPs may have much better effect because it contained much less proteins (∼20% of traditional MPs, Seen in **Figure 2G**).

## Discussion

MPs were first described as subcellular materials originated from platelets in normal plasma and serum^20^. The current diversity of isolated protocols for MPs results in a heterogeneous population of their unknown origins. Moreover, it will be further expanded by the inclusion of additional structures into the pool of MPs, such as apoptotic bodies, migrasomes, or even exosomes^41^. Here we isolated MPs using different centrifugation speed intervals and performed a systematic analysis. We found different methods of isolation of MPs could greatly affect the outcome of MPs enumeration and functional properties, and 3000g is a critical centrifugation speed in determining the MPs composition. Among the particle components obtained under 3000g, platelet-derived particles accounted for more than 80%, while only about 20% of the MPs components obtained by centrifugation above 3000g came from platelets, and the rest came from other cells of the body (**Figure 1**). Therefore, with the significant increase in the proportion of MPs components derived from other cells other than platelets, the characteristic phenotype and function of MPs derived from other cells will undoubtedly be enhanced greatly, and making the functions of MPs more diverse. More importantly, as the proportion of other cell-derived MPs increases, MPs are more likely to become clinical diagnostic markers for some diseases. In addition, the acquisition of plasma MPs is very simple and convenient, and in-depth research will undoubtedly have very important clinical significance, and it can also be an essential research direction for future research on MPs.

Furthermore, the markedly different components of MPs naturally lead to different functions. For example, many previous literatures have reported that procoagulation is a main function of MPs. However, our study found that when most of the small platelets or large platelets fragments were removed by 3000g, the procoagulant function of the MPs was greatly reduced or even disappeared (**Figure 5A**). On the contrary, its functions of promoting angiogenesis and cell migration may be the mainly functions of MPs derived from other cells rather than platelets, because the large platelet-derived particles that were centrifuged at 3000g did not have such functions. This undoubtedly led to a major shift in the understanding of the function of this part of the MPs and clarifies previous misconceptions.

In addition, due to the advantages of simple and convenient acquisition of MPs, making it a biomarker for the diagnosis and prognosis of many major diseases has always been the research goal for relevant research scientists. However, MPs derived from platelets account for 70-90% when isolated with traditional methods (1500g-post), which makes the proportion of MPs from other cells very small proportion and difficult to be detected. Therefore, it has been hard to find the changes in MPs as diagnostic markers for the development of diseases in the past decades. Herein, the platelet-derived MPs in the plasma decreased to about 20%, and the MPs from other cells increased by more than 5 folds with our modified protocol. And our research also showed that the new MPs can be served as a very good marker for the development and treatment of cardiovascular diseases. We are also working on its diagnostic significance in more diseases such as cancer and inflammation. Additionally, our study found that MPs obtained by different centrifugation methods had the function of inhibiting inflammation, so it is speculated that it may be mainly caused by platelet-derived particles. Our research showed that many functions of MPs may need to be re-validated, and clarify which cell-derived MPs may be mediated, so as to conduct in-depth research on its mechanism. This may also be another important direction for MPs studies in the future.

## Methods

### Blood samples

The preparation of microparticle was performed with healthy controls. Written informed consent was obtained from healthy donors under a study protocol approved by the local Institutional Review Board. Citrated blood (4 mL) was collected in an atraumatic fashion using a 21-gauge needle. The samples were processed within two hours.

### The isolation of circulating MPs

Circulating MPs were obtained with three different centrifugation protocols as shown in **Figure 2A**. The centrifugation protocol 1 including two steps was performed according to the traditional protocol, which was centrifuged at 1500g for 20 min and then 20,000g for 30 min to precipitate MPs at 4°C; The protocol 2 include: 1500g was the same to remove cells and platelets, and then centrifuged at 3000g to remove debris of cells, followed by 20000g for 30 mins to get the MPs pellet; The protocol 3 followed as below: 1500, 3000, 5000, 8000 and 12000g for 20 min respectively, finally at 20000g for 20 min to pellet MPs. The pellets containing MPs were fixed or suspended in Phosphate buffer saline (PBS) and stored at −80°C.

### Flow cytometry detection

The effect of centrifugation on detection and quantification of MPs was analyzed by flow cytometry. The microparticle gate was set according to the size and were defined from 50 nm to 1000 nm in diameter. Flow cytometry was performed with a 12-color, triple laser BD FACSLyric flow cytometry (BD Life Sciences, CA, 95131, USA) using BD FACSuite acquisition and analysis software to analyze data. MPs were isolated at different centrifugations and then fixed/permeabilized with BD Cytofix/Cytoperm (BD) for 30 min at 4°C, then washed for 3 times with BD Perm/Wash (BD).

After blocking with Hu Fc Block (BD PMG), samples were incubated with antibodies against CD41a, CD45, CD235a, CD144 and COX4 for 30min, then washed with BD Perm/Wash. The initial gating strategy for MPs is generally based on size by calibration beads. Beads of approximates 1 μm are used to set the upper size limit for MPs detection. Surface markers were normalized using isotype controls. All flow cytometric data were analyzed in Flowjo software.

### Immunofluorescent staining

MPs isolated in different centrifugation were fixed with 4% paraformaldehyde for 30 minutes, then washed for 3 times in PBS. After blocking with 5% BSA, samples were stained with antibodies against CD41, CD45 and Glycophorin at 4°C overnight. Secondary antibody staining was done in blocking buffer at room temperature for 2 h. Samples were imaged using an LSM710 microscope (Zeiss) equipped with a polychromatic META detector.

### Electron microscopy of microparticle samples

Transmission Electron Microscopy (TEM) was used to visualize and measure sizes of MPs. In brief, 5 µL from each sample fraction was placed on a 200-mesh formvar and carbon coated copper grid for 3 min. The excess solution was soaked off by a filter paper before the grid was stained with 5 μl of phosphotungstic acid for 1 min. Excess stain was wicked away to yield a dry grid which was prior to image acquisition by TEM (TALOS F200X scanning/transmission electron microscope) operating at 120 kV.

### Dynamic light scattering (DLS)

MPs size was determined using the Zetasizer ZS (Malvern Panalytical, Malvern, UK) by being exposed to monochromatic light from a laser. MPs scatter the light and undergo Brownian motion which can be analyzed to provide the relative distribution from the different size in terms of how many particles are present. Briefly, samples of MPs were diluted in 1ml of PBS and were measured at 25℃. The analyzers presented microparticle size distribution graphically and in tabular form consisting of a mean size (nm), standard deviation (SD) and polydispersity index (PdI).

### Western blotting

The total proteins from MPs were extracted using RIPA lysis buffer containing phosphatase and protease inhibitor. BCA protein assay kit was used to determine the concentrations of extracted proteins. 10% SDS-PAGE gel was prepared for an equal amount of protein from each sample to run and then transferred to polyvinylidene difluoride (PVDF) membranes. PVDF membranes were blocked with 5% skim milk and incubated with primary antibodies (including CD41, CD36, Perilipin1, Glycophorin, CD45, Tom20, NLRP3, IL-1β, Cleaved Caspase9, iNOS, TNF-α, Tubulin, GAPDH and β-actin) at 4 °C overnight, followed by incubation with secondary horseradish peroxidase conjugated antibodies for 1 h at room temperature.

### Cell culture

Mice Immortalized Bone Marrow-Derived Macrophage (iBMDM), Human Umbilical Vein Endothelial Cells (HUVEC) and Hela cells were cultured in Dulbecco’s modified Eagle’s medium (DMEM) supplemented with 10% fetal bovine serum (FBS), 100 μg/mL streptomycin, and 100 U /mL penicillin, maintained at 37 °C in a humidified atmosphere (5% CO_2_).

### Quantitative reverse transcription PCR

To assess the anti-inflammatory effect, iBMDM cells were seeded into 12-well plates, which were stimulated by LPS (100μg/ml) and then incubated with MPs isolated in different centrifugation for 6h. Total RNA of cell was extracted by Trizol reagent (Takara, Japan), and then transcribed into cDNA by Hifair® II 1st Strand cDNA Synthesis Kit (Yeasen, China). Next, qPCR was performed using Hieff qPCR SYBR Green Master Mix (Yeasen) on LightCycler® 480 Instrument II (Roche). The following Q-PCR conditions were used: 40 cycles of denaturation at 95 °C for 10 s, annealing at 60 °C for 30 s. A comparative threshold cycle method was used to analyze the Q-PCR data, where the amount of target was normalized to the endogenous reference of β-actin in each sample. The primer sequences used were as follows:

Beta-Actin: CGTTGACATCCGTAAAGACC, TAGGAGCCAGAGCAGTAATC; IL1-beta: TTCAGGCAGGCAGTATCACTC, GAAGGTCCACGGGAAAGACAC; NLRP3: GCCGTCTACGTCTTCTTCCTTTC, CATCCGCAGCCAGTGAACAGAG;

### Clotting test *in vitro*

Particles isolated from human blood by different centrifugation protocols were added into the Dulbecco’s modified Eagle’s medium (DMEM) supplemented with 10% fetal bovine serum (FBS). Representative optical images were taken after 5 mins.

As for *in vitro* coagulation test, harvested the 3000g, 1500g-post and 3000g-post particles from mouse blood as described as above protocols, respectively. And then Whole blood samples from C57BL/6 mouse were applied for *in vitro* coagulation test. Sample tubes contained 300 μl whole blood was loaded with 50 μl particles, which come from around 1 ml mouse plasma. The tubes were placed inverted to detect coagulation. And the clotting time was recorded.

### Tube formation: extracellular matrix gel *in vitro* angiogenesis assays

HUVEC were used for tube formation assay. Matrigel (Corning, NY, USA) was melted at 4℃ and coated into 6-well plates with 100 µL per well under sterile conditions without introducing air bubbles. Let the plates sit at 37℃ for at least 15 min to allow gelling of the Matrigel. HUVEC (6×10^4^ cells/well) were then seeded into the pre-coated 6-well plate. Add the MPs in a total volume of 20 µL per well at the same time. The tube-like structures are observed under a microscope at 2h after incubation, and pictures can be taken. Total tube length and individual branch point number were assessed with ImageJ software.

### Transwell assay

In this assay, about 5×10^4^ HUVEC cells in 300 μl FBS-free medium was seeded on top of the filter membrane in a transwell, then inserted into the outer compartment of each well, which contain 750 μl medium with MPs, and incubated for 16 h. fixed and then stained with Hematoxylin and eosin. Pictures were captured and assessed with ImageJ software.

### Anti-inflammatory effect of MPs in LPS-induced acute lung injury *in vivo*

C57BL/6 mice (male, eight weeks) were obtained from the Animal Center of Shanghai Jiao Tong University School of Medicine. The experimental protocol was approved by the Animal Care Committees of Shanghai Jiao Tong University School of Medicine. Mice were anaesthetized with 4% Chloral Hydrate, and 1 mg ‘ultrapure’ LPS (E. coli O111:B4) in 100 μl was instilled intratracheally. Citrated blood was collected from LPS treated mice after 24 h. MPs were isolated using three different centrifugation protocols as described before. And then MPs were washed three times to remove remaining stimulatory factors. Washed MPs were then instilled intratracheally into the new batch of LPS-induced acute lung injury mice. 24h hours after instillation, mice were euthanized, and bronchoalveolar lavage fluid (BALF) samples were analyzed for total protein levels using Bradford assay.

### Statistical analysis

Statistical analyses were conducted using Sigma Plot version 14.0 and GraphPad Prism 9.0 (GraphPad, La Jolla, CA, USA). Quantitative variables are expressed as mean ± SD or median, whereas categorical variables are expressed as numbers and percentages. The Mann-Whitney U-test and ANOVA were used to compare the differences in two or more groups when appropriate. P < 0.05 was considered statistically significant.

## Acknowledgments

This work was supported by the National Natural Science Foundation of China (81770845 and 91854124, To Guorui Huang; 82088102, To Guang Ning; 82171976, To Jiajia Hu) and Shanghai Science & Technology Foundation (18140902800, To Guorui Huang), Shanghai Pujiang Program (21PJD042, To Jiajia Hu).

## References

1. Veziroglu EM and Mias GI. Characterizing Extracellular Vesicles and Their Diverse RNA Contents. Front Genet. 2020;11:700.

2. van Niel G, D’Angelo G and Raposo G. Shedding light on the cell biology of extracellular vesicles. NatRev Mol Cell Biol. 2018;19:213–228.

3. Stahl PD and Raposo G. Extracellular Vesicles: Exosomes and Microvesicles, Integrators of Homeostasis. Physiology (Bethesda). 2019;34:169–177.

4. Yanez-Mo M, Siljander PR, Andreu Z, Zavec AB, Borras FE, Buzas EI, Buzas K, Casal E, Cappello F, Carvalho J, Colas E, Cordeiro-da Silva A, Fais S, Falcon-Perez JM, Ghobrial IM, Giebel B, Gimona M, Graner M, Gursel I, Gursel M, Heegaard NH, Hendrix A, Kierulf P, Kokubun K, Kosanovic M, Kralj-Iglic V, Kramer-Albers EM, Laitinen S, Lasser C, Lener T, Ligeti E, Line A, Lipps G, Llorente A, Lotvall J, Mancek-Keber M, Marcilla A, Mittelbrunn M, Nazarenko I, Nolte-’t Hoen EN, Nyman TA, O’Driscoll L, Olivan M, Oliveira C, Pallinger E, Del Portillo HA, Reventos J, Rigau M, Rohde E, Sammar M, Sanchez-Madrid F, Santarem N, Schallmoser K, Ostenfeld MS, Stoorvogel W, Stukelj R, Van der Grein SG, Vasconcelos MH, Wauben MH and De Wever O. Biological properties of extracellular vesicles and their physiological functions. J Extracell Vesicles. 2015;4:27066.

5. Lacroix R and Dignat-George F. Microparticles as a circulating source of procoagulant and fibrinolytic activities in the circulation. Thromb Res. 2012;129 Suppl 2:S27–9.

6. Sheu JJ, Lee FY, Wallace CG, Tsai TH, Leu S, Chen YL, Chai HT, Lu HI, Sun CK and Yip HK. Administered circulating microparticles derived from lung cancer patients markedly improved angiogenesis, blood flow and ischemic recovery in rat critical limb ischemia. J Transl Med. 2015;13:59.

7. Edelstein LC. The role of platelet microvesicles in intercellular communication. Platelets. 2017;28:222–227.

8. Wan C, Sun Y, Tian Y, Lu L, Dai X, Meng J, Huang J, He Q, Wu B, Zhang Z, Jiang K, Hu D, Wu G, Lovell JF, Jin H and Yang K. Irradiated tumor cell-derived microparticles mediate tumor eradication via cell killing and immune reprogramming. Sci Adv. 2020;6:eaay9789.

9. Jy W, Horstman LL, Jimenez JJ, Ahn YS, Biro E, Nieuwland R, Sturk A, Dignat-George F, Sabatier F, Camoin-Jau L, Sampol J, Hugel B, Zobairi F, Freyssinet JM, Nomura S, Shet AS, Key NS and Hebbel RP. Measuring circulating cell-derived microparticles. J Thromb Haemost. 2004;2:1842–51.

10. Xu R, Greening DW, Zhu HJ, Takahashi N and Simpson RJ. Extracellular vesicle isolation and characterization: toward clinical application. JClinInvest. 2016;126:1152–62.

11. Veerman RE, Teeuwen L, Czarnewski P, Gucluler Akpinar G, Sandberg A, Cao X, Pernemalm M, Orre LM, Gabrielsson S and Eldh M. Molecular evaluation of five different isolation methods for extracellular vesicles reveals different clinical applicability and subcellular origin. J Extracell Vesicles. 2021;10:e12128.

12. Lacroix R, Judicone C, Poncelet P, Robert S, Arnaud L, Sampol J and Dignat-George F. Impact of pre-analytical parameters on the measurement of circulating microparticles: towards standardization of protocol. J Thromb Haemost. 2012;10:437–46.

13. Chandler WL, Yeung W and Tait JF. A new microparticle size calibration standard for use in measuring smaller microparticles using a new flow cytometer. J Thromb Haemost. 2011;9:1216–24.

14. Burnouf T, Goubran HA, Chou ML, Devos D and Radosevic M. Platelet microparticles: detection and assessment of their paradoxical functional roles in disease and regenerative medicine. Blood Rev. 2014;28:155–66.

15. Dey-Hazra E, Hertel B, Kirsch T, Woywodt A, Lovric S, Haller H, Haubitz M and Erdbruegger U. Detection of circulating microparticles by flow cytometry: influence of centrifugation, filtration of buffer, and freezing. Vasc Health Risk Manag. 2010;6:1125–33.

16. Troxler M, Dickinson K and Homer-Vanniasinkam S. Platelet function and antiplatelet therapy. Br J Surg. 2007;94:674–82.

17. Koupenova M, Clancy L, Corkrey HA and Freedman JE. Circulating Platelets as Mediators of Immunity, Inflammation, and Thrombosis. Circ Res. 2018;122:337–351.

18. Tao SC, Guo SC and Zhang CQ. Platelet-derived Extracellular Vesicles: An Emerging Therapeutic Approach. Int J Biol Sci. 2017;13:828–834.

19. Melki I, Tessandier N, Zufferey A and Boilard E. Platelet microvesicles in health and disease. Platelets. 2017;28:214–221.

20. Wolf P. The nature and significance of platelet products in human plasma. Br J Haematol. 1967;13:269–88.

21. Siljander PR. Platelet-derived microparticles - an updated perspective. Thromb Res. 2011;127 Suppl 2:S30–3.

22. Biro E, Sturk-Maquelin KN, Vogel GM, Meuleman DG, Smit MJ, Hack CE, Sturk A and Nieuwland R. Human cell-derived microparticles promote thrombus formation in vivo in a tissue factor-dependent manner. J Thromb Haemost. 2003;1:2561–8.

23. Sims PJ, Faioni EM, Wiedmer T and Shattil SJ. Complement proteins C5b-9 cause release of membrane vesicles from the platelet surface that are enriched in the membrane receptor for coagulation factor Va and express prothrombinase activity. J Biol Chem. 1988;263:18205–12.

24. Sarlon-Bartoli G, Bennis Y, Lacroix R, Piercecchi-Marti MD, Bartoli MA, Arnaud L, Mancini J, Boudes A, Sarlon E, Thevenin B, Leroyer AS, Squarcioni C, Magnan PE, Dignat-George F and Sabatier F. Plasmatic level of leukocyte-derived microparticles is associated with unstable plaque in asymptomatic patients with high-grade carotid stenosis. J Am Coll Cardiol. 2013;62:1436–41.

25. McVey MJ, Spring CM, Semple JW, Maishan M and Kuebler WM. Microparticles as biomarkers of lung disease: enumeration in biological fluids using lipid bilayer microspheres. Am J Physiol Lung Cell Mol Physiol. 2016;310:L802–14.

26. Souza AC, Yuen PS and Star RA. Microparticles: markers and mediators of sepsis-induced microvascular dysfunction, immunosuppression, and AKI. Kidney Int. 2015;87:1100–8.

27. McCarthy EM, Moreno-Martinez D, Wilkinson FL, McHugh NJ, Bruce IN, Pauling JD, Alexander MY and Parker B. Microparticle subpopulations are potential markers of disease progression and vascular dysfunction across a spectrum of connective tissue disease. BBA Clin. 2017;7:16–22.

28. Hough KP, Trevor JL, Strenkowski JG, Wang Y, Chacko BK, Tousif S, Chanda D, Steele C, Antony VB, Dokland T, Ouyang X, Zhang J, Duncan SR, Thannickal VJ, Darley-Usmar VM and Deshane JS. Exosomal transfer of mitochondria from airway myeloid-derived regulatory cells to T cells. Redox Biol. 2018;18:54–64.

29. Stephens OR, Grant D, Frimel M, Wanner N, Yin M, Willard B, Erzurum SC and Asosingh K. Characterization and origins of cell-free mitochondria in healthy murine and human blood. Mitochondrion. 2020;54:102–112.

30. Zhang H, Yu Y, Zhou L, Ma J, Tang K, Xu P, Ji T, Liang X, Lv J, Dong W, Zhang T, Chen D, Xie J, Liu Y and Huang B. Circulating Tumor Microparticles Promote Lung Metastasis by Reprogramming Inflammatory and Mechanical Niches via a Macrophage-Dependent Pathway. Cancer Immunol Res. 2018;6:1046–1056.

31. Schubert P, Culibrk L, Culibrk B, Conway EM, Goodrich RP and Devine DV. Releasates of riboflavin/UV-treated platelets: Microvesicles suppress cytokine-mediated endothelial cell migration/proliferation. Transfusion. 2021;61:1551–1561.

32. Koganti S, Eleftheriou D, Gurung R, Hong Y, Brogan P and Rakhit RD. Persistent circulating platelet and endothelial derived microparticle signature may explain on-going pro-thrombogenicity after acute coronary syndrome. Thromb Res. 2021;206:60–65.

33. Li X, Tse HF and Jin LJ. Novel endothelial biomarkers: implications for periodontal disease and CVD. J Dent Res. 2011;90:1062–9.

34. Koga H, Sugiyama S, Kugiyama K, Watanabe K, Fukushima H, Tanaka T, Sakamoto T, Yoshimura M, Jinnouchi H and Ogawa H. Elevated levels of VE-cadherin-positive endothelial microparticles in patients with type 2 diabetes mellitus and coronary artery disease. J Am Coll Cardiol. 2005;45:1622–30.

35. Bulut D, Maier K, Bulut-Streich N, Borgel J, Hanefeld C and Mugge A. Circulating endothelial microparticles correlate inversely with endothelial function in patients with ischemic left ventricular dysfunction. J Card Fail. 2008;14:336–40.

36. Jung C, Sorensson P, Saleh N, Arheden H, Ryden L and Pernow J. Circulating endothelial and platelet derived microparticles reflect the size of myocardium at risk in patients with ST-elevation myocardial infarction. Atherosclerosis. 2012;221:226–31.

37. Mezentsev A, Merks RM, O’Riordan E, Chen J, Mendelev N, Goligorsky MS and Brodsky SV. Endothelial microparticles affect angiogenesis in vitro: role of oxidative stress. Am J Physiol Heart Circ Physiol. 2005;289:H1106–14.

38. Liang HZ, Li SF, Zhang F, Wu MY, Li CL, Song JX, Lee C and Chen H. Effect of Endothelial Microparticles Induced by Hypoxia on Migration and Angiogenesis of Human Umbilical Vein Endothelial Cells by Delivering MicroRNA-19b. Chin Med J (Engl). 2018;131:2726–2733.

39. Garnier Y, Ferdinand S, Garnier M, Cita KC, Hierso R, Claes A, Connes P, Hardy-Dessources MD, Lapoumeroulie C, Lemonne N, Etienne-Julan M, El Nemer W and Romana M. Plasma microparticles of sickle patients during crisis or taking hydroxyurea modify endothelium inflammatory properties. Blood. 2020;136:247–256.

40. Zhang D, Lee H, Wang X, Groot M, Sharma L, Dela Cruz CS and Jin Y. A potential role of microvesicle-containing miR-223/142 in lung inflammation. Thorax. 2019;74:865–874.

41. Ma L, Li Y, Peng J, Wu D, Zhao X, Cui Y, Chen L, Yan X, Du Y and Yu L. Discovery of the migrasome, an organelle mediating release of cytoplasmic contents during cell migration. Cell Res. 2015;25:24–38.

